# Recurrent evolution of cross-resistance in response to selection for improved post-infection survival in *Drosophila melanogaster*

**DOI:** 10.1101/2021.11.26.470139

**Authors:** Aparajita Singh, Aabeer Basu, Biswajit Shit, Tejashwini Hegde, Nitin Bansal, Nagaraj Guru Prasad

## Abstract

The host susceptibility to one pathogen can decrease, increase, or remain unaffected by virtue of the host evolving resistance towards a second pathogen. Negative correlations between a host susceptibility to different pathogens is an often-cited explanation for maintenance of genetic variation in immune function determining traits in a host population. In this study, we investigated the change in susceptibility of *Drosophila melanogaster* flies to various novel bacterial pathogens after being experimentally selected for increased resistance to one particular bacterial pathogen. We independently selected flies to become more resistant towards *Enterococcus faecalis* and *Pseudomonas entomophila*, and baring a few exceptions the evolved populations exhibited cross-resistance against the range of pathogens tested in the study. Neither the identity of the native pathogen nor the host sex was major determining factors in predicting the pattern of cross-resistance exhibited by the selected populations. We therefore report that a generalized cross-resistance to novel pathogens can repeatedly evolve in response to selection for resistance against a single pathogen.

## Introduction

Omnipresence of parasites and pathogens poses a continuous threat to host fitness by compromising host life-span, fecundity, or both. Evolving defence mechanisms against parasites and pathogens is therefore of great importance to the hosts. Continuous selection for better defence should erode additive genetic variation for defence related traits, as more and more resistant genotypes are driven to fixation (Schelunburg et al. 2009, Lazzaro and Little 2009). Empirical studies have repeatedly found evidence contradicting this theoretical expectation. Genetic variation for anti-pathogen defence have been reported in studies, both in field and lab, across various organisms: (viz. Tinsley et al. 2006, Lazzaro et al. 2006, Raberg et al. 2007). Various factors can contribute towards this difference between the predicted and observed results, including cost and condition-dependence of immune defence, host-pathogen co-evolution and resultant evolution of specific defence, and variation in biotic and abiotic environment (Schmid-Hempel 2003, Lazzaro and Little 2009).

Hosts and pathogens exist as part of a complex network of interactions, and hosts are rarely challenged by a single pathogen in the wild (Betts et al. 2016). Under such circumstances where a host must counter multiple pathogens, hosts can evolve a generic defence mechanism to counter all threats, or different host genotypes may specialise against different types of pathogens, leading to evolution of immune specificity (Decaestecker et al. 2003, Schmid-Hempel and Ebert, 2003). Increased resistance against one pathogen can produce corelated decrease (positive cross-resistance) or increase (negative cross-resistance) in susceptibility towards a second pathogen (Fellowes et al. 1999, Kraaijeveld et al. 2012).

At the phenotypic level, cross-resistance manifests when hosts infected with one pathogen show increased or decreased susceptibility to a second pathogen. For example, mice infected with *Schistosomatium douthitti* are more resistant to subsequent infection by *Schistosoma mansoni* (Hunter et al. 1961). Similarly, *Anopheles gambiae* mosquito hosts that are previously exposed to *Vavria culicis* are more resistant to *Plasmodium berghei* (Bargielowski and Koella 2009). In *Drosophila melanogaster* flies, infection with any one of *Providencia rettgeri*, *Enterococcus faecalis* and *Serratia marcescens* make flies more resistance towards later infection by the other two pathogens (Chambers et al. 2019).

At evolutionary level, cross-resistance is determined by how a host evolved to counter a particular pathogen responds to infection by a novel pathogen. Iso-female lines of *Drosophila melanogaster* show positive correlation for resistance to two parasitoids *Leptopilina boulardi* and *Leptopilina heterotoma* (Boulétreau and Wajnberg 1986; Delpuech et al. 1994). *D. melanogaster* populations selected for increased resistance to *L. boulardi* had increased resistance to *Asobara tabida* and *L. heterotoma* where as those populations selected for increased resistance to *A. tabida* had higher resistance to *L. heterotom*a (Fellows et al. 1999). Martins et al (2013) selected *D. melanogaster* populations for increased survivorship against infection from *Pseudomonas entomophila* and found that the evolved populations were also better at surviving *Pseudomonas putida*. *D. melanogaster* populations selected against DCV showed positive cross-resistance against Cricket Paralysis Virus (CrPV) and Flock House Virus (FHV) (Martins et al. 2014). In contrast to these results of the evolution of positive cross-resistance, many other studies have found no evidence for the evolution of cross-resistance as a result of evolution towards a particular pathogen/parasite. Martins et al. (2013) found that their populations selected for increased survivorship against *P. entomophila* was as good as the controls in surviving infections from *Erwinia carotovora* or *Serratia marcescens*. Selection for increased resistance against the parasitoid *A. tabida* did not increase resistance to the parasitoid *L. boulardi* (Fellows et al. 1999), the microsporidian *Tubulinosema kingi* or the fungus *Beauveria bassiana* (Kraaijeveld et al. 2012). Populations of *D. melanogaster* evolved against *Bacillus cereus* did not evolve cross-resistance to DSV (Bentz et al. 2017). Similarly, greater wax moth evolved against *B. bassiana* did not evolve resistance to *Metarhizium anisopliae* (Dubovskiy et al. 2013). *Tribolium castaneum* larvae coevolved with *B. bassiana* was cross resistant to *Bacillus thuringiensis* but not *P. entomophila* (Biswas et al. 2018). To the best of our knowledge, only one study, has found the evolution of negative cross-resistance. Martins et al. (2013) found that populations of *D. melanogaster* evolved against *P. entomophila* were more susceptible to infections from *Enterococcus faecalis*, DCV and FHV. To summarise, at the evolutionary level, most of the studies have either found evidence for positive cross-resistance or have found no evidence for cross-resistance. There is little empirical evidence for the evolution of negative cross-resistance.

The variation in outcomes of the above described studies may be attributed to numerous factors: (a) the phylogenetic relatedness between the pathogen used for experimental evolution and the pathogens used for testing cross-resistance (Schmid-Hempel and Ebert 2003), (b) common mechanisms of pathogen virulence or host resistance (Vallet-Gely et al. 2008, Dubovskiy et al. 2013), (c) route of infection (Martins et al. 2013, Biswas et al. 2018), and (d) the genetic architecture of the host-population in the study. Additionally, host-sex is a major determinant of host immune function. The sexes differ from one another in terms of optimal life-history, environmental infection risk, and physiological modulators of immunity (viz. testosterone in male mammals and juvenile hormone in female insects); these factors together contribute towards sexual dimorphism in immune function (Zuk and McKean 1996 , Rolff et al. 2002, Schmid-Hempel and Ebert 2003, Nunn et al. 2009, Vincent and Sharp 2014, Sharp and Vincent 2015), and can potentially be another factor that leads to differential patterns of cross-resistance observed in empirical studies. Studies on immunity often focus on only one sex, and even when both sexes are used in experiments, the statistical analysis does not involve sex as a factor. Hence, sexual dimorphism in cross-resistance has not been explored to a great extent in the existing literature.

In the present study we created two selection regimes using replicate populations of *Drosophila melanogaster* that share a close common ancestry (figure 1). We then selected one set of populations for increased survival post systemic infection with a Gram-negative bacterial pathogen, *Pseudomonas entomophila* (Gupta et al 2016). The other set of populations were similarly selected against systemic infection from a Gram-positive bacterium, *Enterococcus faecalis*. In both the selection regimes, populations quickly evolved improved survival against infection from the native pathogen, exhibiting increased post-infection survival compared to paired controls (Gupta et al. 2016, data presented here). Using these two sets of populations we tested (a) if adapting to one pathogen confers the hosts cross-resistance to novel pathogens; (b) is the pattern of cross-resistance contingent on the identity of the native pathogen; and (c) is cross-resistance sexually dimorphic? We use the phrase *pathogen resistance* to imply the ability of the host to survive a challenge with a pathogen, unless mentioned otherwise.

**Figure 1.**
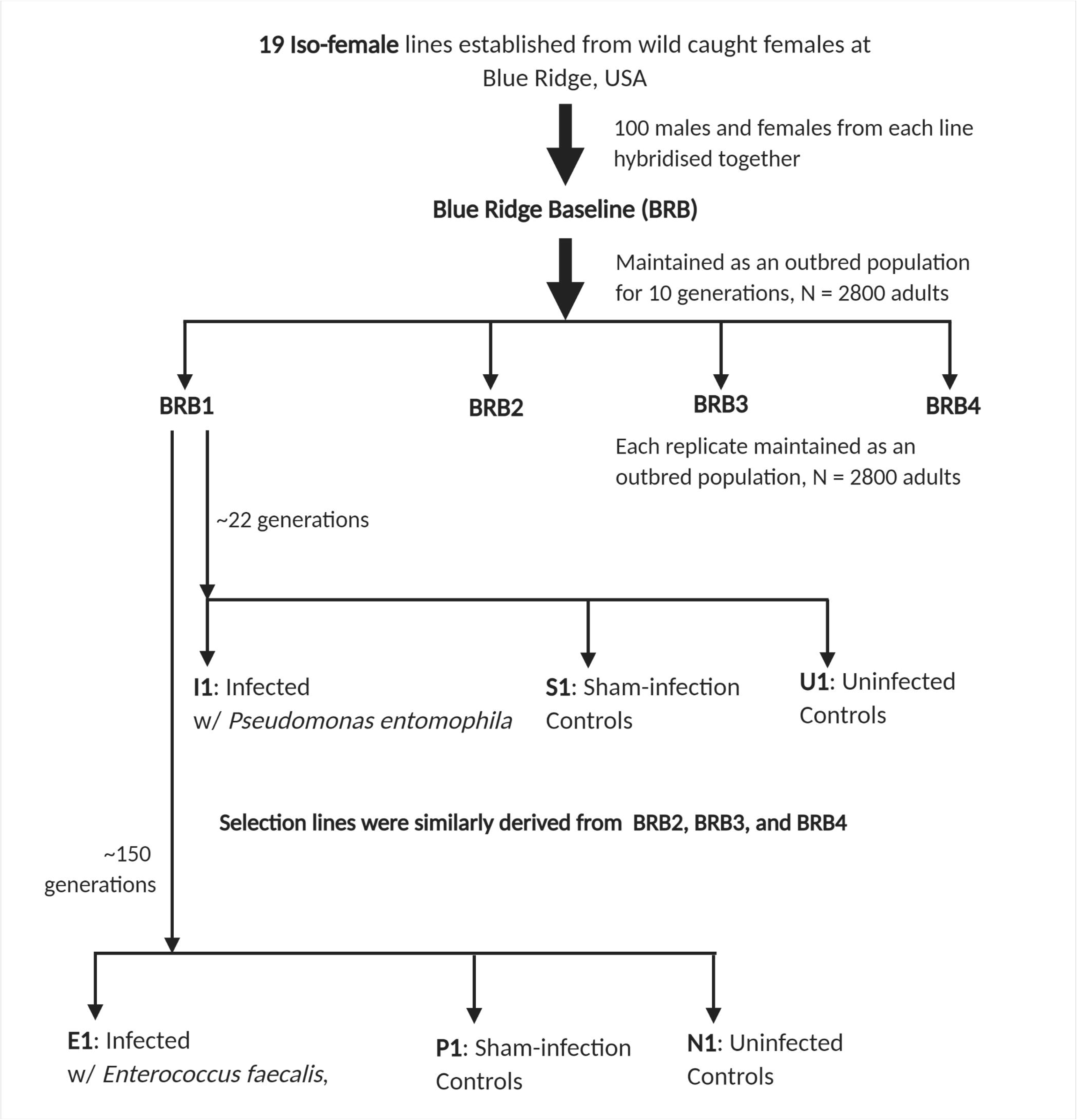
The ancestry of the populations used in this study: inter-relatedness of BRB populations and the two selection lineages (EPN and IUS populations).

## Materials and methods

For the present study we used two sets of selected populations of *Drosophila melanogaster*, each selected for improved post-infection survival when infected with a different entomopathogenic bacteria.

### Ancestral populations

Both sets of selected populations were derived from a common ancestor, the Blue Ridge Baseline (BRB), a wild-type, outbred population, with four evolutionary replicates, BRB1-4. The derivation and maintenance protocol of the BRB populations are described in detail in Singh et al. (2015). Briefly, each replicate is maintained at a census size of 2800 adults, on a 14-day discrete generation cycle, at 25 °C on a 12:12 light-dark cycle and 50-60% relative humidity on standard banana-yeast-jaggery food. Juveniles of these populations are reared in 40 glass vials (25mm diameter × 90mm height) at a density of ∼70 eggs per vial, with 6-8 mL of standard banana-jaggery-yeast medium. By 12th day post egg laying (PEL), almost all flies eclose and the adults are transferred to Plexiglass cages (25 cm length x 20 cm width x 15 cm height) having food in petri plate (90 mm diameter) supplemented with live yeast paste. On the 14th day PEL, fresh food plates (cut into halves to expose the vertical surfaces where the flies seem to prefer to lay eggs, hereafter called as cut-plate) are provided in the cages for 18 hours of egg laying. Eggs are collected from these cut-plates at above mentioned densities and dispensed into glass vials to start a new generation.

### Selection for increased survival against systemic infection with *Enterococcus faecalis* (*Ef*): EPN populations

Three populations were derived from each replicate population of BRB after 150 generations: (a) E1, infected with *<u>E</u>nterococcus faecalis*, (b) P1, <u>p</u>ricking control, and (c) N1, <u>n</u>ormal control were derived from BRB1. Similarly, we derived E2, P2 and N2 from BRB2 ancestor, and so on. Therefore, there were totally 12 populations in this selection regime: E1-4, P1-4, and N1-4. Populations with the same numeral shared a more recent common ancestor. For example, E1, P1 and N1 were more closely related to each other than any of them is to E2, P2, N2 etc. Additionally, populations bearing the same numeral were always handled together, during selection and during experimentation. Therefore, populations with the same numeral were treated as statistical blocks. Consequently, we had four blocks (Block 1-4) in the EPN selection regime (E1, P1, N1 forming block 1 and so on). For all populations, eggs were collected at a density of 60-80 eggs per vial (25 mm diameter × 90 mm height) containing 6-8 ml of food (similar to the ancestral population) in 10 such vials and were incubated at standard laboratory conditions as mentioned above. By 9th-10th day 95% of the flies eclose. Further handling depended on the type of selection being imposed.

In the E1-4 populations, on day 12 PEL, when the flies are 2-3 days old as adults, from each of the 10 juvenile development vials, we randomly chose 20 females and 20 males flies, and infected them with the pathogen by septic injury on the thorax with a Minutien pin (0.1 mm, Fine Science Tools, USA) dipped in a bacterial suspension (in MgSO_4_ saline buffer) at optical density (OD_600_) of 0.8, under light CO_2_ anaesthesia. Therefore, a total of 200 females and 200 males are infected every generation for each E population. After infections the flies were shifted to a Plexiglas cage (14 cm length x 16 cm width x 13 cm height) provided with a food plate (60 mm petri plate in diameter); fresh food plates were provided every alternate day. For flies infected with *Enterococcus faecalis* majority of the mortality is observed between 18 and 48 hours post-infection with very few flies dying before 18 or after 48 hours. After 96 hours post-infection, fifty percent of the infected flies in each E populations would survive to contribute to the next generation. Throughout the selection history of these populations, the pathogen infection dose was modulated to induce fifty percent mortality: this ensured a constant, directional selection process. Therefore, flies of zeroth generation were infected with OD_600_=0.8, and the infection dose was increased to OD_600_=1.0 at generation 23, and again increased to OD_600_=1.2 at generation 41. 96 hours after infection (day 16 PEL) the population cages are provided with fresh oviposition plates (cut-plate) and 18 hours later eggs were collected of these plates to start the next generation.

Flies of the P1-4 populations are maintained identically to the E populations, except that (a) on day 12 PEL, when the flies are 2-3 days old as adults, they are pricked with a Minutien pin (0.1mm, Fine Science Tools, USA) dipped in sterile MgSO_4_ buffer under light CO_2_ anaesthesia, before being placed in cages; (b) From each of the 10 juvenile development vials, we randomly chose 10 females and 10 males such that 100 females and 100 males are sham-infected every generation for each population. There is negligible mortality (1-2%) in these cages between the time of infection and oviposition.

Flies of the N1-4 are maintained identical to P populations except that on day 12 PEL we randomly chose 10 females and 10 males from each of the 10 juvenile development vials under CO_2_ anaesthesia such that 100 females and 100 males are subjected to uninfected treatment every generation for each population. There is negligible mortality in these cages.

Each block was handled on a different day, i.e., E1, P1, and N1 were handled together on one day; E2, P2, and N2 were handled together on the next day, and so on. Every generation, in each population the number of eclosing flies is the same (about 700). The flies eclose in the vials by day 10 and would have mated by day 12. On day 12, we subsample 400 flies (200 of each sex) in each of the E populations and subject them to infections. Of these about 100-120 flies per sex survive till day 16 and contribute to the next generation. From the P and N populations, on day 12, we subsample 200 flies (100 of each sex). There is negligible mortality (1-2%) in the P and N regimes. Therefore, on day 16, when we collect eggs to start the next generation, close to 100 flies of each sex are present in these populations. Thus, our protocol ensures that the number of adults at the time of egg collection are similar across populations. The EPN selection regime is thus maintained on a 16-day discrete generation cycle.

### Selection for increased survival against systemic infection with *Pseudomonas entomophila* (*Pe*): IUS populations

The IUS populations were similarly derived from the BRB populations after 22 generations of establishment of the base populations, and have been previously described in Gupta et al. (2016). Briefly, three selection regimes were derived from each replicate population of BRB: (a) I1-4, infected with *Pseudomonas entomophila*, (b) S1-4, <u>s</u>ham-infected control, and (c) U1-4, <u>u</u>ninfected controls were derived from BRB1, and so on. The maintenance of these lines is identical to that of the EPN lines, except that (a) I,U,S populations were started from BRB populations after 22 generations of lab adaptation while E,P,N populations were started from BRB populations after 150 generations of lab adaptation, (b) in the I1-4 populations 150 females and 150 males are infected every generation for each population whereas in E1-4 200 females and males, (c) in I,U,S Gram-negative bacteria *Pseudomonas entomophila* and E,P,N Gram-positive *Enterococcus faecalis* is used, (d) peak mortality window for I is 20 hours to 60 hours and for E is 18 hours to 48 hours. The derivation of both selection regime is summarized in figure 1 and Table 4.

### Bacterial stocks and infection procedure

The bacterial stocks were maintained as 17% glycerol stocks frozen at -80 °C. To obtain fresh bacterial cells for infection (either for regular selection protocol or during experimental infections), 10 ml lysogeny broth (Luria-Bertani-Miller, HiMedia) is inoculated with a stab of bacterial glycerol stock and incubated overnight at appropriate temperature with continuous mixing at 150 RPM. A secondary culture is established by inoculating 10 ml lysogeny broth using 100 ul of the overnight culture; this secondary culture is incubated at appropriate temperature till desired turbidity is reached. The bacterial cells from this culture is pelleted down via centrifugation and resuspended in sterile MgSO_4_ buffer (10 mM) at the required optical density (OD_600_). Flies are infected by pricking them in the thorax under light CO_2_ anaesthesia with a 0.1Minutein pin (Fine Scientific Tolls, USA) dipped in the bacterial suspension. Sham infection are carried out similarly except with a pin dipped in sterile MgSO_4_.

Seven pathogens were used in total in this study. Four Gram-positive bacteria: *Enterococcus faecalis* (hereafter *Ef*, grown at 37°C), *Bacillus thuringiensis* (hereafter *Bt*, grown at 30° ), *Micrococcus luteus* (hereafter *Ml*, grown at 37°C), and *Staphylococcus succinus* (hereafter *Ss*, grown at 37°C) were used. Three Gram-negative bacteria: *Pseudomonas entomophila* (hereafter *Pe*, grown at 27°C), *Erwinia c. carotovora* (hereafter *Ecc*, grown at 30°C), and *Providencia rettgeri* (hereafter *Pr*, grown at 37°C) were used.

### Pre-experiment standardization

To account for any non-genetic parental effects, experimental eggs were collected from flies which were grown in common garden conditions for one generation (Rose, Evolution 1984). Eggs were collected from all the populations at a density of 60-80 eggs per vial: 10 such vials were established per population. The eggs completed their development into adults in these vials, and on day 12 PEL, the adults were transferred to plexiglass cages (14×16×13 cm^3^) with food plates (petri plates, 60 mm diameter). Eggs for experimental flies were collected from these population cages.

### Rearing of the Experimental Flies

Three days prior to the egg collection, food plates supplemented with live yeast were provided to the standardised flies in the cages. After 2 days, yeast plate was replaced with cut-plate for the next 18 hours for egg laying. From these cut-plates eggs were transferred into glass vials (25 vials per population to test for response to selection and 50 vials per population to test for cross-resistance), at the density of 60-80 eggs per vial (90 mm x 25 mm), each vial having 6-8ml of standard banana-jaggery food. The vials were incubated under conditions identical to the maintenance of the selection regime. Eggs developed into adults in these vials within 10 days after egg collection, and the adults remained in these vials till day 12 PEL, wherefrom they were used for experiments.

### Test of response to selection in EPN populations

This experiment was carried out after 35 generations of forward selection. On 12^th^ day PEL, we sampled 400 females and 400 males from each of the E, P, and N populations. These were then randomly assigned to one of the three treatments: (a) Infection treatment: 200 females and 200 males were infected with *Enterococcus faecalis* (*Ef)* at OD_600_ = 0.8; (b) Sham-infection treatment: 100 females and 100 males were sham-infected with sterile MgSO_4_ solution; and (c) Uninfected treatment: 100 females and 100 males were subjected to light CO_2_ anaesthesia only. Post treatment, the flies were placed inside plexiglass cages (14×16×13 cm^3^) containing food in petri plates (60 mm diameter). Mortality was noted every 4-6 hours until 96 hours post infection for each cage. Fresh food plates were provided to the cages on every alternate day. Individual blocks were handled on separate days. Altogether, 200 flies/sex/population/block were infected with *Ef*, 100 flies/sex/population/block were sham-infected, and 100 flies/sex/population/block were kept as uninfected control.

### Test of response to selection in IUS populations

This experiment was carried out after 160 generations of forward selection. On 12^th^ day PEL, we sampled 100 females and 100 males from each of the I, U, and S populations. These were then randomly assigned to one of the two treatments: (a) Infection treatment: 50 females and 50 males were infected with *Pseudomonas entomophila* (*Pe*) at OD_600_ = 1.5; and (b) Sham-infection treatment: 50 females and 50 males were sham-infected with sterile MgSO_4_ solution. Post treatment, the flies were placed inside plexiglass cages (14×16×13 cm^3^) containing food in petri plates (60 mm diameter). Mortality was noted every 4-6 hours until 96 hours post infection for each cage. Fresh food plates were provided to the cages on every alternate day. Individual blocks were handled on separate days. Altogether, 50 flies/sex/population/block were infected with *Pe*, and 50 flies/sex/population/block were used as sham-infected controls.

### Test of cross-resistance in EPN populations

Test for cross-resistance in the EPN populations were carried out after 40 generations of forward selection. Flies from E and P populations were infected with 6 pathogens: *Bacillus thuringiensis* (*Bt*), *Micrococcus luteus* (*Ml*), *Staphylococcus succinus* (*Ss*), *Erwinia c. carotovora* (*Ecc*), *Pseudomonas entomophila* (*Pe*), and *Providencia rettgeri* (*Pr*) along with sham-infected controls. Infection dose for all pathogens was OD_600_ = 1.0. On 12^th^ day PEL (flies were 2-3 day old as adults), 50 flies/sex/pathogen/population were infected and transferred to plexiglass cages (14 x 16 x 13 cm^3^) with food plates (60 mm diameter). Mortality was noted every 4-6 hours until 96 hours post infection for each cage. Fresh food plates were provided to the cages on every alternate day. Individual blocks of the selection regime were handled on different days.

### Test of cross-resistance in IUS populations

Test for cross-resistance in the IUS populations were carried out after 160 generations of forward selection. Flies from I and S populations were infected with 6 pathogens: *Bacillus thuringiensis* (*Bt*), *Micrococcus luteus* (*Ml*), *Staphylococcus succinus* (*Ss*), *Enterococcus faecalis* (*Ef*), *Erwinia c. carotovora* (*Ecc*), and *Providencia rettgeri* (*Pr*) along with sham-infected controls. Infection dose for all pathogens was OD_600_ = 1.0. On 12^th^ day PEL (flies were 2-3 day old as adults), 50 flies/sex/pathogen/population were infected and transferred to plexiglass cages (14 x 16 x 13 cm^3^) with food plates (60 mm diameter). Mortality was noted every 4-6 hours until 96 hours post infection for each cage. Fresh food plates were provided to the cages on every alternate day. Individual blocks of the selection regime were handled on different days.

### Statistical analysis

All analyses was performed using R statistical software, version 4.1.0 (R Core Team 2021). Mixed-effect cox-proportional hazard models were fitted to the data using the *coxme* function of the “coxme” package (Therneau 2020), and the confidence intervals for these models were calculated using *confint* function of the base R package. Survival curves were plotted using the *ggsurvplot* function of the “survminer” (Kassambara et al. 2021) package after modelling the data using *survfit* function from the “survival” (Therneau 2021) package.

For the analysis of data from the response to selection experiments we first modelled the total data as:

survival ∼ infection treatment + (1|block),

where infection treatment was considered as a fixed factor and block as a random factor. Next, we modelled the data from only the infected individuals as:

survival ∼ selection regime + sex + (selection regime : sex) + (1|block),

where selection regime, sex and their interaction were considered as fixed factors and block as random factor. This was done separately for the EPN and the IUS selection regimes.

For the analysis of the data from the cross-resistance experiments we modelled the data for each pathogen as:

survival ∼ selection regime + sex + (selection regime : sex) + (1|block),

where selection regime, sex and their interaction were considered as fixed factors and block as random factor. This was done separately for the EPN and the IUS selection regimes.

## Results

### Test of response to selection in EPN populations, selected for resistance against *Enterococcus faecalis*

The EPN selection regime consists of three types of populations: (a) E1-4: selected for increased resistance against *Enterococcus faecalis*; (b) P1-4: pricking (sham-infected) controls; and (c) N1-4: normal (uninfected) controls (See ‘Materials and Methods’ for more details). To test for the primary response to selection, we infected the E, P, and N populations with *Enterococcus faecalis* (*Ef*) at OD_600_ = 0.8 after 35 generations of forward selection.

Both the infected and the uninfected treatments are significantly different from sham-infected treatment (table 1), infected treatment surviving less (hazard ratio 14.6184, 95% CI 11.8732, 17.9980) and uninfected treatment surviving more (hazard ratio 0.0851, 95% CI 0.0413, 0.1753) than sham-control treatment. Comparing within the infected treatment, E populations, survived significantly better than flies from the P control populations (hazard ratio 0.636, 95% CIs 0.5454, 0.7417; figure 2); the N and P populations were similar in post-infection survival (hazard ratio 1.0464, 95% CI 0.9103, 1.2028). Males were not significantly different from females in terms of post-infection survival (hazard ratio 0.9203, 95% CI 0.7980, 1.0614).

**Figure 2.**
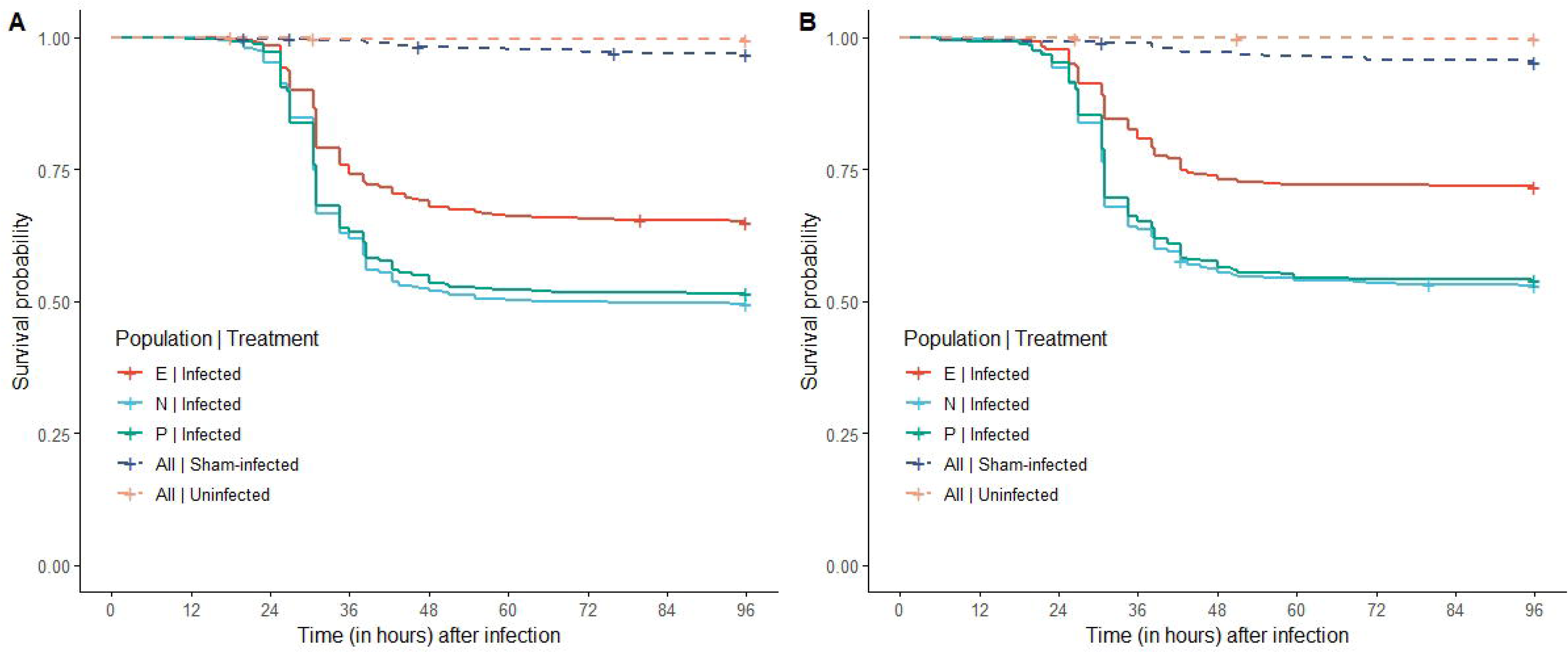
Survival curves for (A) females and (B) males of EPN selection regime tested for response to selection after 35 generations of forward selection, infected with their primary pathogen *Enterococcus faecalis*.

**Table 1.** Output of mixed-effects Cox proportional hazards analysis of flies of EPN (generation 35) and IUS (generation 160) selection regimes tested for response to selection by being infected with their respective primary pathogens, *Enterococcus faecalis* and *Pseudomonas entomophila*. Hazard ratios are relative to the default level for each factor, which is set at 1. The default level for “Treatment” is ‘Sham-infected’, the default level for “Selection” is ‘P’ or ‘S’ depending upon the regime under analysis, and the default level for “Sex” is ‘Females’. Hazard ratio greater than 1 implies reduced survival compared to the default level. Significant effects are marked in bold.

|  | Hazards Ratio | Lower CI (95%) | Upper CI (95%) | Z | p-value | Variance (for random factor) |
| --- | --- | --- | --- | --- | --- | --- |
| A. Effect of infection treatments on overall survival in EPN selection regime. |  |  |  |  |  |  |
| Treatment *Uninfected* | **0.0851** | **0.0413** | **0.1753** | **-6.69** | **2.3 e-11** |  |
| Treatment *Infected* | **14.6184** | **11.8732** | **17.9980** | **25.28** | **0.0 e+00** |  |
| Block |  |  |  |  |  | 0.1008 |
| B. Effect of selection history and sex on survival of infected flies in EPN selection regime. |  |  |  |  |  |  |
| Selection *E* | **0.6360** | **0.5454** | **0.7417** | **-5.77** | **7.9 e-09** |  |
| Selection *N* | 1.0464 | 0.9103 | 1.2028 | 0.64 | 5.2 e-01 |  |
| Sex *Males* | 0.9203 | 0.7980 | 1.0614 | -1.14 | 2.5 e-01 |  |
| Selection *E* : Sex *Males* | 0.8337 | 0.6651 | 1.0450 | -1.58 | 1.1 e-01 |  |
| Selection *N* : Sex *Males* | 0.9935 | 0.8133 | 1.2137 | -0.06 | 9.5 e-01 |  |
| Block |  |  |  |  |  | 0.1142 |
| C. Effect of infection treatments on overall survival in IUS selection regime. |  |  |  |  |  |  |
| Treatment *Infected* | **14.7962** | **11.1879** | **19.5685** | **18.89** | **0.0 e+00** |  |
| Block |  |  |  |  |  | 0.1060 |
| D. Effect of selection history and sex on survival of infected flies in IUS selection regime. |  |  |  |  |  |  |
| Selection *I* | **0.0944** | **0.0568** | **0.1568** | **-9.12** | **0.0 e+00** |  |
| Selection *U* | 0.8942 | 0.6919 | 1.1556 | -0.85 | 3.9 e-01 |  |
| Sex *Males* | **2.1630** | **1.7063** | **2.7420** | **6.38** | **1.8 e-10** |  |
| Selection *I* : Sex *Males* | 0.8899 | 0.4691 | 1.6883 | -0.36 | 7.2 e-01 |  |
| Selection *U* : Sex *Males* | 0.7332 | 0.5192 | 1.0354 | -1.76 | 7.8 e-02 |  |
| Block |  |  |  |  |  | 0.2160 |

### Test of response to selection in IUS populations, selected for resistance against *Pseudomonas entomophila*

The IUS selection regime consists of three types of populations: (a) I1-4: selected for increased resistance against *Pseudomonas entomophila*; (b) S1-4: sham-infected controls; and (c) U1-4: uninfected controls (See ‘Materials and Methods’ for more details). To test for the primary response to selection, we infected the IUS populations with *Pseudomonas entomophila* (*Pe*) at OD_600_ = 1.5 after 160 generations of forward selection.

Infected flies (hazard ratio 14.7962, 95% CI 11.1879, 19.5685) survive significantly less compared to flies from sham-infected treatment (table 1). I populations, when infected, survived significantly better than flies from the S control populations (hazard ratio 0.0944, 95% CIs 0.0568, 0.1568) (figure 3). However, the two control populations, U and S, were similar in terms of post-infection survival (hazard ratio 0.8942, 95% CIs 0.6919, 1.1556). Male survived significantly less when infected compared to females (hazard ratio 2.1630, 95% CIs 1.7063, 2.7420).

**Figure 3.**
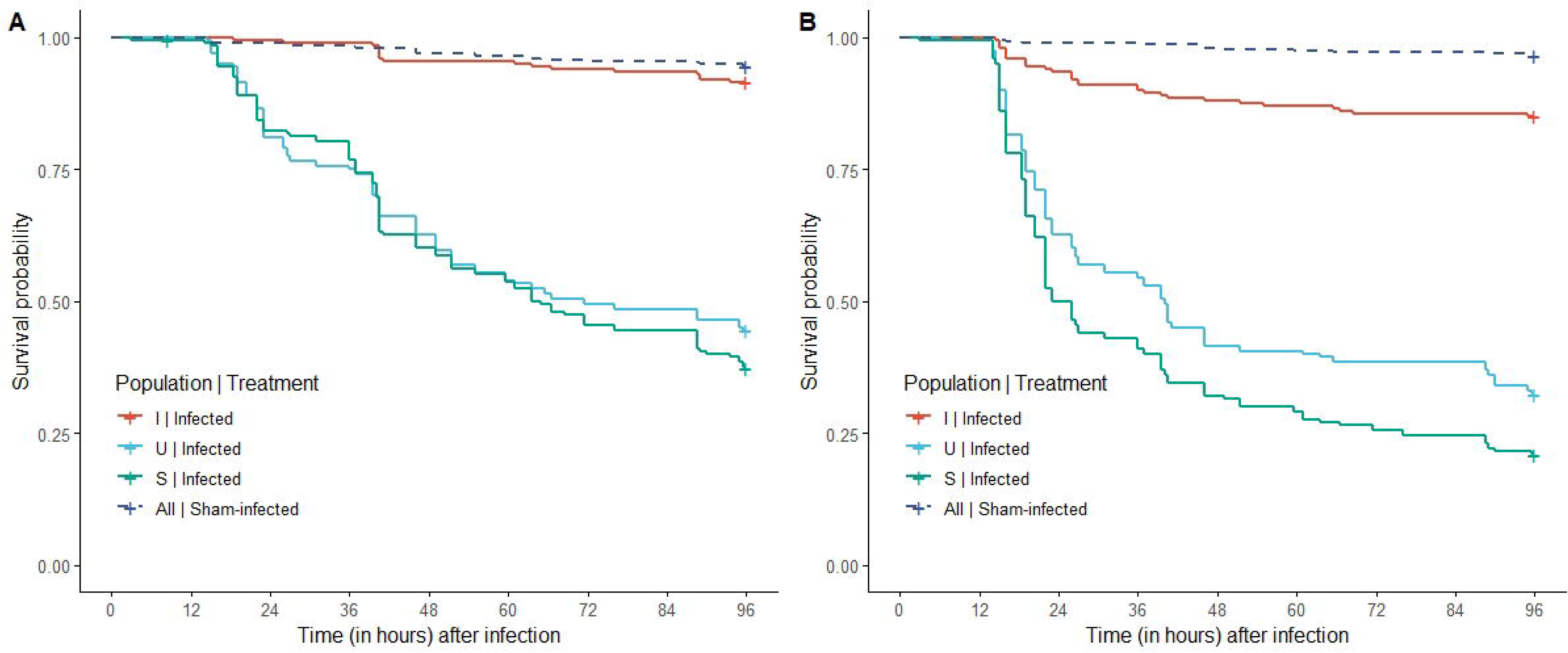
Survival curves for (A) females and (B) males of IUS selection regime tested for response to selection after 160 generations of forward selection, infected with their primary pathogen *Pseudomonas entomophila*.

### Test of cross-resistance against novel pathogens in EPN populations, selected for resistance against *Enterococcus faecalis*

Test for cross-resistance in the EPN populations, after 40 generations of forward selection, flies from E (selected) and P (pricking controls) populations were infected with six novel pathogens: *Bacillus thuringiensis* (*Bt*), *Micrococcus luteus* (*Ml*), *Staphylococcus succinus* (*Ss*), *Erwinia c. carotovora* (*Ecc*), *Pseudomonas entomophila* (*Pe*), and *Providencia rettgeri* (*Pr*), (along with sham-infected controls) with infection dose for all pathogens maintained at OD_600_ = 1.0.

E populations are significantly better in post-infection survival from P populations when infected with five out of six novel pathogens: *Bt* (hazard ratio 0.7108, 95% CIs 0.5682, 0.8892), *Ml* (hazard ratio 0.7099, 95% CIs 0.5206, 0.9681), *Ss* (hazard ratio 0.5158, 95% CIs 0.3497, 0.7609), *Ecc* (hazard ratio 0.6409, 95% CIs 0.4802, 0.8556), and *Pe* (hazard ratio 0.4604, 95% CIs 0.3501, 0.6054). The E and P populations were not significantly different in terms of survival when infected with *Pr* (hazard ratio 0.7585, 95% CIs 0.5459, 1.0538). Males and females did not differ from one another in post-infection survival when infected with any of the six pathogens (figure 4, table 2).

**Figure 4.**
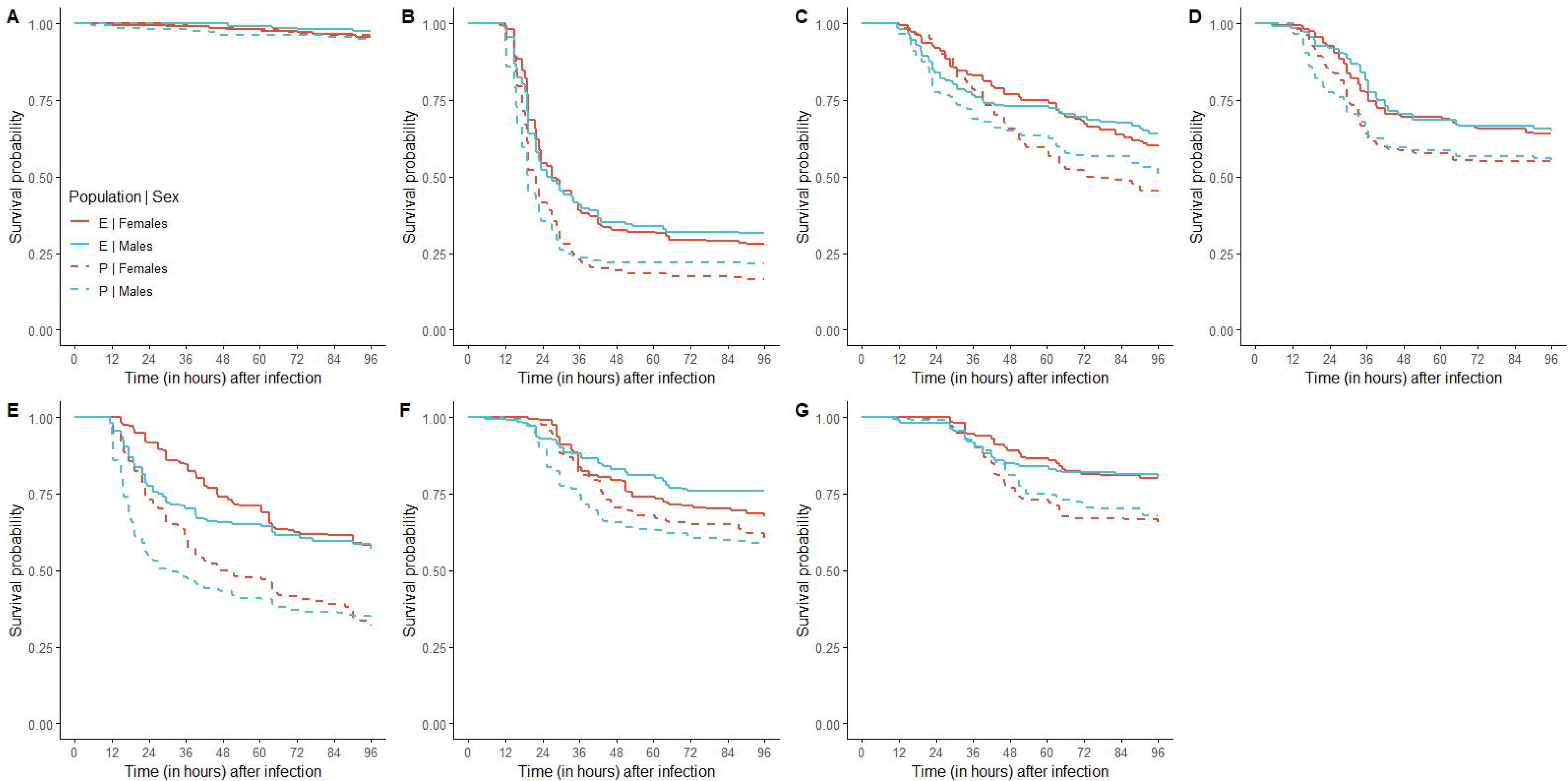
Survival curves for flies of EPN selection regime (only E and P populations were used in this experiment) tested for cross-resistance against novel pathogens after 40 generations of forward selection: (A) Sham-infection controls, (B) *Bacillus thuringiensis*, (C) *Erwinia c. carotovora*, (D) *Micrococcus luteus*, (E) *Pseudomonas entomophila*, (F) *Providencia rettgeri*, and (G) *Staphylococcus succinus*.

**Table 2.** Output of mixed-effects Cox proportional hazards analysis of flies of EPN selection regime (generation 40; only E and P populations were used in this experiment) infected with novel pathogens to test for cross-resistance. Hazard ratios are relative to the default level for each factor, which is set at 1. The default level for “Selection” is ‘P’ and the default level for “Sex” is ‘Females’. Hazard ratio greater than 1 implies reduced survival compared to the default level. Significant effects are marked in bold. (Pathogens marked with ‘#’ are of the same Gram-character as the primary pathogen used in the selection regime.)

|  | Hazards Ratio | Lower CI (95%) | Upper CI (95%) | Z | p-value | Variance (for random factor) |
| --- | --- | --- | --- | --- | --- | --- |
| A. Bacteria: *Bacillus thuringiensis* (Bt)<sup>#</sup>. |  |  |  |  |  |  |
| Selection *E* | **0.7108** | **0.5682** | **0.8892** | -<br>**2.99** | **0.0028** |  |
| Sex *Males* | 1.0593 | 0.8518 | 1.3174 | 0.52 | 0.6000 |  |
| Selection *E* : Sex *Males* | 0.9242 | 0.6713 | 1.2725 | -<br>0.48 | 0.6300 |  |
| Block |  |  |  |  |  | 0.1360 |
| B. Bacteria: *Erwinia c. carotovora* (Ecc). |  |  |  |  |  |  |
| Selection *E* | **0.6409** | **0.4802** | **0.8556** | -<br>**0.32** | **0.0025** |  |
| Sex *Males* | 0.9258 | 0.7048 | 1.2159 | -<br>0.55 | 0.5800 |  |
| Selection *E* : Sex *Males* | 1.0079 | 0.6629 | 1.5326 | 0.04 | 0.9700 |  |
| Block |  |  |  |  |  | 0.0572 |
| C. Bacteria: *Micrococcus luteus* (Ml)<sup>#</sup>. |  |  |  |  |  |  |
| Selection *E* | **0.7099** | **0.5206** | **0.9681** | -<br>**2.16** | **0.03** |  |
| Sex *Males* | 1.0152 | 0.7573 | 1.3609 | 0.10 | 0.92 |  |
| Selection *E* : Sex *Males* | 0.9387 | 0.6042 | 1.4585 | -<br>0.28 | 0.78 |  |
| Block |  |  |  |  |  | 0.0436 |
| D. Bacteria: *Pseudomonas entomophila* (Pe). |  |  |  |  |  |  |
| Selection *E* | **0.4604** | **0.3501** | **0.6054** | -<br>**5.55** | **2.8 e-08** |  |
| Sex *Males* | 1.1667 | 0.9176 | 1.4834 | 1.26 | 2.1 e-01 |  |
| Selection *E* : Sex *Males* | 0.9950 | 0.6769 | 1.4625 | -<br>0.03 | 9.8 e-01 |  |
| Block |  |  |  |  |  | 0.1214 |
| E. Bacteria: *Providencia rettgeri* (Pr). |  |  |  |  |  |  |
| Selection *E* | 0.7585 | 0.5459 | 1.0538 | -<br>1.65 | 0.099 |  |
| Sex *Males* | 1.1193 | 0.8215 | 1.5252 | 0.71 | 0.480 |  |
| Selection *E* : Sex *Males* | 0.6553 | 0.4036 | 1.0642 | -<br>1.71 | 0.088 |  |
| Block |  |  |  |  |  | 0.1046 |
| F. Bacteria: *Staphylococcus succinus* (Ss)<sup>#</sup>. |  |  |  |  |  |  |
| Selection *E* | **0.5158** | **0.3497** | **0.7609** | -<br>**3.34** | **0.0008** |  |
| Sex *Males* | 0.8880 | 0.6327 | 1.2464 | -<br>0.69 | 0.4900 |  |
| Selection *E* : Sex *Males* | 1.1356 | 0.6510 | 1.9808 | 0.45 | 0.6500 |  |
| Block |  |  |  |  |  | 0.0417 |

### Test of cross-resistance against novel pathogens in IUS populations, selected for resistance against *Pseudomonas entomophila*

Test for cross-resistance in the IUS populations, after 160 generations of forward selection, flies from I (selected) and S (sham-infection controls) populations were infected with six novel pathogens: *Bacillus thuringiensis* (*Bt*), *Micrococcus luteus* (*Ml*), *Staphylococcus succinus* (*Ss*), *Enterococcus faecalis* (*Ef*), *Erwinia c. carotovora* (*Ecc*), and *Providencia rettgeri* (*Pr*), (along with sham-infected controls) with infection dose for all pathogens maintained at OD_600_ = 1.0.

I populations are significantly better in post-infection survival from S populations when infected with five out of six novel pathogens: *Ef* (hazard ratio 0.7238, 95% CIs 0.5376, 0.9746), *Ml* (hazard ratio 0.6562, 95% CIs 0.4804, 0.8962), *Ss* (hazard ratio 0.5002, 95% CIs 0.3366, 0.7433), *Ecc* (hazard ratio 0.1072, 95% CIs 0.0661, 0.1738) and *Pr* (hazard ratio 0.2473, 95% CIs 0.1522, 0.4020). The I and S populations were not significantly different in terms of survival when infected with *Bt* (hazard ratio 1.1228, 95% CIs 0.8779, 1.4360). Additionally, males survived significantly less than females when infected with two out of six novel pathogens: *Ecc* (hazard ratio 1.8834, 95% CIs 1.4825, 2.3927) and *Pr* (hazard ratio 1.8229, 95% CIs 1.3492, 2.4630), but males and females were equally susceptible to the rest of the four pathogens (figure 5, table 3).

**Figure 5.**
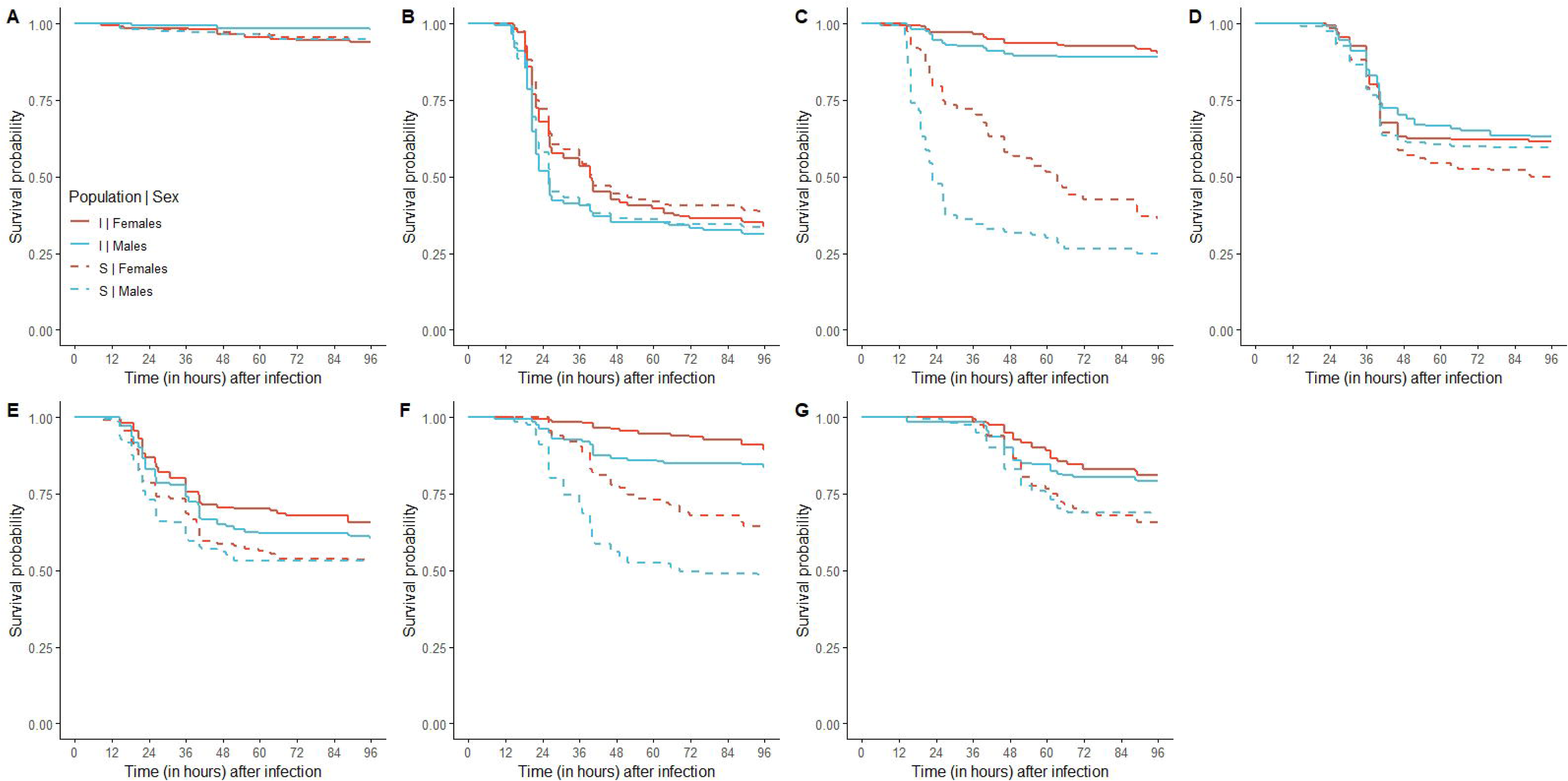
Survival curves for flies of IUS selection regime (only I and S populations were used in this experiment) tested for cross-resistance against novel pathogens after 160 generations of forward selection: (A) Sham-infection controls, (B) *Bacillus thuringiensis*, (C) *Erwinia c. carotovora*, (D) *Enterococcus faecalis*, (E) *Micrococcus luteus*, (F) *Providencia rettgeri*, and (G) *Staphylococcus succinus*.

**Table 3.** Output of mixed-effects Cox proportional hazards analysis of flies of IUS selection regime (generation 160; only I and S populations were used in this experiment) infected with novel pathogens to test for cross-resistance. Hazard ratios are relative to the default level for each factor, which is set at 1. The default level for “Selection” is ‘S’ and the default level for “Sex” is ‘Females’. Hazard ratio greater than 1 implies reduced survival compared to the default level. Significant effects are marked in bold. (Pathogens marked with ‘#’ are of the same Gram-character as the primary pathogen used in the selection regime.)

|  | Hazards Ratio | Lower CI (95%) | Upper CI (95%) | Z | p-value | Variance (for random factor) |
| --- | --- | --- | --- | --- | --- | --- |
| A. Bacteria: *Bacillus thuringiensis* (Bt). |  |  |  |  |  |  |
| Selection I | 1.1228 | 0.8779 | 1.4360 | 0.92 | 0.360 |  |
| Sex Males | 1.2728 | 0.9953 | 1.6275 | 1.92 | 0.055 |  |
| Selection I : Sex Males | 1.0005 | 0.7101 | 1.4098 | 0.00 | 1.000 |  |
| Block |  |  |  |  |  | 0.1288 |
| B. Bacteria: *Erwinia c. carotovora* (Ecc)<sup>#</sup>. |  |  |  |  |  |  |
| Selection I | **0.1072** | **0.0661** | **0.1738** | -<br>**9.06** | **0.0 e+00** |  |
| Sex Males | **1.8834** | **1.4825** | **2.3927** | **5.18** | **2.2 e-07** |  |
| Selection I : Sex Males | 0.6341 | 0.3282 | 1.2254 | -<br>1.36 | 1.8 e-01 |  |
| Block |  |  |  |  |  | 0.1204 |
| C. Bacteria: *Enterococcus faecalis* (Ef). |  |  |  |  |  |  |
| Selection I | **0.7238** | **0.5376** | **0.9746** | -<br>**2.13** | **0.033** |  |
| Sex Males | 0.8115 | 0.6054 | 1.0879 | -<br>1.40 | 0.160 |  |
| Selection I : Sex Males | 1.1394 | 0.7388 | 1.7573 | 0.59 | 0.550 |  |
| Block |  |  |  |  |  | 0.0378 |
| D. Bacteria: *Micrococcus luteus* (Ml). |  |  |  |  |  |  |
| Selection I | **0.6562** | **0.4804** | **0.8962** | -<br>**2.65** | **0.0081** |  |
| Sex Males | 1.0676 | 0.8014 | 1.4222 | 0.45 | 0.6500 |  |
| Selection I : Sex Males | 1.1253 | 0.7306 | 1.7334 | 0.54 | 0.5900 |  |
| Block |  |  |  |  |  | 0.0891 |
| E. Bacteria: *Providencia rettgeri* (Pr)<sup>#</sup>. |  |  |  |  |  |  |
| Selection I | **0.2473** | **0.1522** | **0.4020** | -<br>**5.64** | **1.7 e-08** |  |
| Sex Males | **1.8229** | **1.3492** | **2.4630** | **3.91** | **9.2 e-05** |  |
| Selection I : Sex Males | 0.9194 | 0.4924 | 1.7167 | -<br>0.26 | 7.9 e-01 |  |
| Block |  |  |  |  |  | 0.0452 |
| F. Bacteria: *Staphylococcus succinus* (Ss). |  |  |  |  |  |  |
| Selection I | **0.5002** | **0.3366** | **0.7433** | -<br>**3.43** | **0.0006** |  |
| Sex Males | 0.9396 | 0.6677 | 1.3221 | -<br>0.36 | 0.7200 |  |
| Selection I : Sex Males | 1.2306 | 0.7057 | 2.1459 | 0.73 | 0.4600 |  |
| Block |  |  |  |  |  | 8.399 e-05 |

**Table 4.** Comparison of the maintenance regime of the ancestral and the selected populations.

|  | BRB | EPN | IUS |
| --- | --- | --- | --- |
| **Generation cycle** | 14-days | 16-days | 16-days |
| **Food medium** | Banana-jaggery-yeast medium | Banana-jaggery-yeast medium | Banana-jaggery-yeast medium |
| **Egg density per vial** | 60-80 eggs | 60-80 eggs | 60-80 eggs |
| **Number of vials per population** | 40 | 10 for E, 10 for P, and 10 for N | 10 for I, 10 for S, and 10 for U |
| **Number of replicate populations** | 4 (BRB1-4) | 12 (E1-4, P1-4, and N1-4) | 12 (I1-4, S1-4, and U1-4) |
| **Yeast prior to egg collection** | Yes | No | No |
| **Pathogen used** | NA | *Enterococcus faecalis* (Ef), Gram-positive bacteria | *Pseudomonas entomophila* (Pe), Gram-negative bacteria |
| **Ancestral population** | NA | BRB1-4 | BRB1-4 |
| **Number of generations after which populations were started from ancestral population** | NA | 150 generations | 22 generations |
| **Number of adults in each generation (during the window of reproduction)** | 2800 adults, sex ratio not maintained artificially | Approximately 100 females and 100 males per population | Approximately 100 females and 100 males per population |
| **Peak mortality window** | NA | Between 18 hours and 48 hours for E populations | Between 20 hours and 60 hours for I populations |

## Discussion

Hosts selected to be more immune-competent against one particular pathogen can evolve correlated resistance to other pathogens. We tested for evolution of such cross-resistance in two sets of replicate *Drosophila melanogaster* populations, one selected for resistance against a Gram-negative bacterial pathogen *Pseudomonas entomophila* (Gupta et al. 2016) for 160 generations, and another selected for resistance against a Gram-positive pathogen *Enterococcus faecalis* (reported here for the first time) for 40 generations. Each selected regime and its corresponding paired controls were infected with six novel bacterial pathogens to test if the selected populations were better at surviving a pathogen challenge compared to the controls. Our primary observations from these experiments are as follows:

a. Evolution of increased immunity: The E populations, that were selected for resistance against *Enterococcus faecalis* showed rapid evolution to the selection pressure. After 35 generations of selection, post-infection survival of the E (selected) populations was better than both P (sham-infected control) and N (un-infected control) populations (figure 2, Table 1) when infected with *E. faecalis*. Host sex didn’t have a significant effect on post-infection survival (Table 1). Similarly, the I populations, selected for resistance against *Pseudomonas entomophila*, were better at surviving a challenge with *P. entomophila* compared to both S (sham-infected control) and U (un-infected control) populations (figure 3, Table 1), after 160 generations of selection. Post-infection survival of females was better than males for all three populations (I, U, and S; Table 1).
b. Evolution of cross-resistance: When challenged with six novel pathogens, the E (selected) populations were less susceptible to infections, compared to the P (control) populations, to all the novel bacteria except *Providencia rettgeri*, for which there was no difference in the post-infection survival of E and P populations (figure 4, Table 2). Similarly, the I (selected) populations survived better compared to the S (control) populations when challenged with six novel pathogens, except for *Bacillus thuringiensis*, for which I and S populations exhibited equal mortality (figure 5, table 3). Therefore, out of twelve total tests for cross-resistance (two selection lines × six novel pathogens) we found evidence for positive cross-resistance in ten comparisons and no effect of selection in the remaining two. We did not observe a single case of negative cross-resistance. Interestingly the population selected against *E. faecalis* were resistant to *P. entomophila* and the populations selected against *P. entomophila* were resistant to *E. faecalis*.
c. Sexual dimorphism: For the populations selected against *E. faecalis* sex had no effect on post-infection survival when the populations were challenged either by the native pathogen or by the six novel pathogens (figure 4, Table 2). For populations selected against *P. entomophila* females exhibited reduced mortality compared to males for all Gram-negative pathogens (native and novel), but not in case of the Gram-positive pathogens (all novel) (figure 5, Table 3). Sex-by-population interaction was not observed for any bacteria for either of the two selection regimes.

In case of both of the selected populations we noted the evolution of cross-resistance against a wide range of pathogens. Previous studies, using similar selection designs have reported evolution of cross-resistance against only a few limited pathogens. For example, Martins et al. (2013) reported that flies evolved to be resistant against *P. entomophila* were cross-resistant against *P. putida* only, and exhibited increased or no change in susceptibility when infected with *E. faecalis*, and *S. marcescens* and *E. carotovora*, respectively; the selected flies were also more susceptible than controls when infected with viruses. For neither of our selected populations did we observe a scenario where the selected populations were more susceptible to a novel pathogen compared to controls. More importantly, our populations evolved to resist *P. entomophila* were also resistant to *E. faecalis*; another point of difference between our results and that reported by Martins et al. (2013).

There can be two possible explanations for the different outcomes of the two studies. One, the genetic architecture of the starting base-line populations is a major determinant of the outcome of any selection experiment. The genotypic co-variances of susceptibilities to different pathogens in our starting populations might have been different from that of Martin et al. (2013). Two, correlated responses to selection observed upon in selection experiments can depend upon the number of generations of forward selection (Chippindale et al.1997, Tetonio and Rose 2000). Martins et al. (2013) tested for cross-resistance after 27-30 generations of forward selection whereas we tested for cross-resistance after 160 (IUS populations) generations. It is possible that populations subjected to sustained to long term directional selection exhibit a broader range of cross-resistance.

With respect to the predictive effect of the identity of the native pathogen (the pathogen used for selection) on the pattern of cross-resistance exhibited by the selected populations, we expected that the selected populations would be more resistant to novel pathogens that are most similar to the native pathogen phylogenetically and mechanistically (in terms of both pathogen virulence and host resistance). In contradiction with our expectation, both of our selected populations (E and I populations) evolved cross-resistance against a wide range of bacterial pathogens, barring a few exceptions: E populations did not exhibit cross-resistance against *P. rettgeri*, while the I populations did not exhibit cross-resistance against *B. thuringiensis*. In both cases the novel pathogen was on the opposite Gram-character to that of the native pathogen, and hence phylogenetic dissimilarity, or more accurately, cellular/morphological dissimilarity, may be invoked as an explanation. We did not test the phylogenetic dissimilarity hypothesis to its full extent given we used only bacterial pathogens as novel challenges.

Previous studies that have tested evolved flies with different taxa of pathogens/parasites have reported mixed results. Kraaijeveld et al. (2012) selected *D. melanogaster* flies for improved defence against parasitoid *Asobara tabida* and found that selection had no effect on defence against fungal pathogen *Beauveria bassiana* and microsporidian pathogen *Tubulinosema kingi*. In the same study, selection for increased defence against *B. bassiana* had no effect on defence against *A. tabida* (Kraaijeveld et al. 2012). Martins et al. (2013) found that selecting *D. melanogaster* flies for resistance against bacteria *P. entomophila* increases their susceptibility to Drosophila C Virus and Flock House Virus. Biswas et al (2018) selected *Tribolium castaneum* beetles for resistance against fungus *B. bassiana*, and found that defence against bacteria *B. thuringiensis* increased as a consequence of selection, while defence against bacteria *P. entomophila* was compromised.

Alternatively, results from our experiments may indicate shared pathways of pathogen virulence or hosts defence. Insect immunity is a composite trait with multiple layers of complexity. Post-infection survival is a function of the hosts ability to both control the systemic proliferation of pathogens and to deal with the systemic damage incurred in its interaction with the pathogen (Dionne and Schneider 2008, Raeberg et al. 2009). Insects have multiple mechanism of resisting pathogen growth, some specific while some general, with these mechanisms not always acting in a mutually exclusive manner (Lemaitre and Hoffmann 2007). For example, defence against systemic infection by *Enterococcus faecalis* requires two cellular defence mechanisms: phagocytosis (Nehme et al 2011) and melanization (Ayres and Schneider 2008), and also the involvement of genes downstream of the toll signalling pathway (Gobert et al. 2003, Nehme et al 2011, Hanson et al 2019). Cross-resistance can be driven by the overlap of either the virulence traits employed by the pathogen (Vallet-Gely et al. 2008) or the common mechanism of defence utilized by the host. Previous research has suggested that there is some order of specificity of immune defences at the level of the Gram-character of the bacterial pathogens (Lemaitre and Hoffmann 2007). Our results from the EPN selection regime indicate that flies evolved to counter *Enterococcus faecalis* infection were better at defending against novel pathogens independent of the pathogen identity. This points at the fact that the selected populations have evolved a mechanism which can serve as a general defence against a wide variety of novel bacterial pathogen. Phagocytosis or melanization are the most likely candidates given these pathways tend to be more generalist defence mechanisms compared to IMD/Toll regulated anti-microbial peptide (AMP) based humoral defences (Lemaitre and Hoffmann 2007).

There is limited information regarding the mechanism of host defence against systemic infection by *Pseudomonas entomophila*. The primary reason for this is that most studies have focused on elucidating the host’s response to oral infection by *Pseudomonas entomophila* (Vodovar et al. 2005, Liehl et al. 2006), however there is some indication that response to systemic infection and oral infection share certain common mechanisms (Martin et al 2013). Previous research had shown that flies evolved to fight off systemic infection by *Pseudomonas entomophila* are less susceptible to infection by other Gram-negative bacteria compared to controls while having similar or increased susceptibility to Gram-positive bacteria and viruses (Martin et al 2013). This is congruent with the IMD/Toll dichotomy of defence mechanisms. Our results, on the other hand, show that flies evolved to survive systemic challenge with *Pseudomonas entomophila* exhibit reduced susceptibility to a variety of pathogens independent of their identity; with the only exception being *Bacillus thuringiensis*, in the case of which both selected and control populations were equally susceptible. Interestingly, the mechanistic basis of virulence following oral infection by *Pseudomonas entomophila* share certain common features with that of *Bacillus thuringiensis*. Crystal proteins produced by *Bacillus thuringiensis* perforate the gut wall of insects (Bravo et al. 2007, Soberon et al. 2007); a similar role is played by monolysin produced by *Pseudomonas entomophila* (Opota et al. 2011). Resistance against *Pseudomonas entomophila* is driven by genes downstream of the Imd signalling pathway, which includes AMPs such as diptericin, diptericin b, cecropin A1, attacin A, attacin C, cecropin C, drosomysin and drosopterin (Vodovar et al. 2005). These same AMPs are required for defence against both oral and systemic infection by *Pseudomonas entomphila*; the same set of genes is also required for defence against *Erwinia carotovora carotovora* (Vodovar et al. 2005). Given that infection by *Pseudomonas entomophila* induces the expression of a very wide range of AMPs, evolution of cross-resistance against a wide variety of pathogens is not surprising.

Sexual dimorphism in immune function has been theoretically predicted and empirically established by previous studies (Zuk and McKean 1996, Rolff et al. 2002, Schmid-Hempel and Ebert 2003, Nunn et al. 2008, Vincent and Sharp 2014, Sharp and Vincent 2015). In populations selected for defence against *E. faecalis* no effect of sex on post-infection survival was observed, when the selected and the paired control populations were challenged with the native or the novel pathogens. Effect of sex was seen in case of populations selected against *P. entomophila*, but only for Gram-negative pathogens; no sex × selection regime interaction was observed in case of any pathogen. Therefore, although we observed evolution of sexual dimorphism in one of our selection regimes, sexual dimorphism had no deterministic contribution towards the pattern of cross-resistance observed in our experiments.

To summarize, in this paper we report that selecting replicate *D. melanogaster* populations against either *Enterococcus faecalis* or *Pseudomonas entomophila* leads to the correlated evolution of cross-resistance of a wide variety of novel pathogens. The identity of the native pathogen did not predict the novel pathogens against which the selected populations exhibited cross-resistance but it did predict the novel pathogens against which the selected populations did not show cross-resistance. Furthermore, the pattern of cross-resistance observed in case of either of the selected populations were not affected by sex of the host; even in cases where sex affected host infection survival the effects were similar for the selected and the control populations. Differences in susceptibility of a host to different pathogens is one of the common justifications for presence of genetic variation in immune function related traits in natural populations. Our results suggest that hosts can become cross-resistant to a variety of pathogens by virtue of evolving to resist a single pathogen, and therefore positive correlations between host’s resistance against different pathogens may not be very rare in nature. In this study, we tested the evolution of cross-resistance against novel pathogens one at a time. It would be interesting to study how our selected populations fair in comparison to the controls when co-infected with more than one pathogen, because co-infecting pathogens often interact amongst themselves and with the host in unique ways that are not apparent in single infections.

## Author contributions

Fund acquisition: NGP; Conceptualization and study design: AS, AB; Execution of experiments and data collection: AS, AB, BS, TH, NB; Statistical analysis: AS, AB; Manuscript preparation: AS, AB, NGP. All authors have consented to the final form of the manuscript submitted.

## Acknowledgements

This study was funded by IISER Mohali intramural funds and a research grant (no. BT/PR14278/BRB/10/1417/2015) from Dept. of Biotechnology, Govt. of India. AS is supported by Junior and Senior Research Fellowships from University Grant Council, Govt. of India. AB is supported by Junior and Senior Research Fellowships from Council of Scientific and Industrial Research, Govt. of India. BS is supported by INSPIRE Scholarship for Higher Education, and TH and NB are supported by KVPY Fellowship, both from Dept. of Science and Technology, Govt. of India. We thank Prof. P Cornelis (Free University, Brussels, Belgium) for providing us *Pseudomonas entomophila*, Prof. B Lazzaro (Cornell University, Ithaca, USA) for providing us *Enterococcus faecalis* and *Providencia rettgeri*, Dr. E Succena and T Paulo (The Instituto Gulbenkian Ciência, Oeiras, Portugal) for providing us *Erwinia c. carotovora* and Dr. K Singh (IISER Mohali, India) for isolating the *Staphylococcus succinus* bacteria, pathogens used in this study.

